# Rapid and scalable profiling of nascent RNA with fastGRO

**DOI:** 10.1101/2020.01.24.916015

**Authors:** Elisa Barbieri, Connor Hill, Mathieu Quesnel-Vallieres, Yoseph Barash, Alessandro Gardini

## Abstract

Genome-wide profiling of nascent RNA has become a fundamental tool to study transcription regulation. Over the past decade, next-generation sequencing has fostered development of a handful of techniques (i.e. GRO-seq, PRO-seq, TT-seq and NET-seq) that map unprocessed transcripts originating from both the coding and the noncoding portion of the genome. Unlike steady-state RNA sequencing, nascent RNA profiling mirrors the real-time activity of RNA Polymerases and provides an accurate readout of transcriptome-wide variations that occur during short time frames (i.e. response to external stimuli or rapid metabolic changes). Some species of nuclear RNAs, albeit functional, have a short half-life and can only be accurately gauged by nascent RNA techniques (i.e. lincRNAs and eRNAs). Furthermore, these techniques capture uncapped post-cleavage RNA at termination sites or promoter-associated antisense RNAs, providing a unique insight into RNAPII dynamics and processivity.

Here we present a run-on assay with 4s-UTP labelling, followed by reversible biotinylation and affinity purification via streptavidin. Our protocol allows streamlined sample preparation within less than 3 days. We named the technique fastGRO (fast Global Run-On). We show that fastGRO is highly reproducible and yields a more complete and extensive coverage of nascent RNA than comparable techniques. Importantly, we demonstrate that fastGRO is scalable and can be performed with as few as 0.5×10^6 cells.

## Introduction

In slightly more than a decade, NextGen Sequencing technology has revolutionized the field of transcription by allowing precision mapping of most RNA species, from mRNAs to lincRNAs. Usually, RNA is extracted from crude cell extracts via acidic phenol-chloroform precipitation and either reverse transcribed with oligodT or subjected to ribosomal RNA-depletion, first, followed by reverse transcription using a pool of short random oligos (thus avoiding the polyadenylation bias). Adapter ligation and PCR-based amplification convert the original pool of RNAs into a sequencing-ready library that will generate quantitative transcriptomic profiles (Stark, Grzelak et al. 2019). Regardless the specific protocol of choice, there are several limitations to these widely-used RNA-seq techniques. First, traditional RNA-seq measures steady-state, mostly cytoplasmic, RNA species. Steady-state RNA levels are the ultimate result of synthesis rate, RNA processing and RNA stability. Both RNA processing and stability are highly regulated in every cell type (Schoenberg and Maquat 2012; Pai and Luca 2019; Yamada and Akimitsu 2019). Therefore, RNA-seq alone is insufficient to infer the accurate transcriptional activity of any given gene. Additionally, transcription by RNA Polymerase II (RNAPII) is not a steady and passive process of ribonucleotide chain assembly. There are multiple, critical regulatory steps and checkpoints all along the transcription cycle that cannot be discerned with the resolution offered by RNA-seq (Adelman and Lis 2012; Kwak and Lis 2013; Proudfoot 2016). Lastly, there are low-abundant and poorly stable RNA species that fall below the RNA-seq detection threshold. For instance, biologically active enhancer RNAs and other lincRNA species are hardly represented in conventional transcriptomic data (Lai and Shiekhattar 2014; Gardini and Shiekhattar 2015). To overcome limitations of RNA-seq, several groups have developed high-throughput methods that tap into the so-called ‘nascent’ fraction of cellular RNA (Wissink, Vihervaara et al. 2019). Nascent transcripts represent the small RNA fraction (>0.5 % of total RNA content in a cell) that is actively synthesized and still associated with RNA Polymerase. Global Run-On sequencing (GRO-seq) was the first genome-wide technique developed to probe nascent transcription genome-wide (Core, Waterfall et al. 2008). GRO-seq yields an exact footprint of already engaged RNA polymerase, by building upon the strengths of a 40-year old assay (Gariglio, Buss et al. 1974; Gariglio, Bellard et al. 1981). In GRO-seq, nuclei are isolated and flash-frozen, only to resume transcription *in vitro* in the presence of a labelled nucleotide (Core, Waterfall et al. 2008; Gardini 2017). PRO-seq was developed years later as a modification of GRO-seq using biotinylated nucleotides (Core, Martins et al. 2014). Both techniques are time-consuming and marred by non-standardized library preparation(Mahat, Kwak et al. 2016; Gardini 2017). Another popular method for deep sequencing of nascent RNA, TT-seq (and its parent technique 4SU-seq) relies on metabolic labelling of RNA but also recovers partly and fully processed RNA that is not associated with RNAPII (Schwanhausser, Busse et al. 2011; Schwalb, Michel et al. 2016). Additional strategies to purify RNAPII-associated transcripts, such as NET-seq and mNET-seq, are biased towards identifying pausing sites of polymerase and depend on affinity purification or subcellular fractionation (Mayer, di Iulio et al. 2015; Nojima, Gomes et al. 2016).

We have developed a run-on technique (fast Global Run-On, fastGRO) that allows robust mapping of the nascent transcriptome in under 3 days. Our technique optimizes the usage of 4-thio ribonucleotides (4-S-UTP) for nuclear run-on assays. We take advantage of reversible biotinylation to label and enrich for newly synthesized RNA species and we ultimately generate strand-specific libraries for Illumina sequencing using commercially available prep kits. Here, we employ fastGRO to measure nascent RNA in HeLa and THP1 cells and we compare our technique to a variety of nascent RNA assays that have been widely adopted over the past years. While reducing processing time by more than half, we show that fastGRO yields more consistent coverage across gene bodies (spanning introns and post-termination RNAs) than other benchmark techniques. We also find that processed RNA contamination is significantly lower in fastGRO. We use fastGRO to measure a variety RNA species, including low-abundant lincRNAs and short-lived eRNAs and antisense promoter transcripts. A major limitation of current techniques is the large amount of starting material required, which restricts their applicability to inexpensive, fast-growing cell lines. Here we show that fastGRO is down-scalable and can be performed with as few as 0.5-1x 10^6 cells, potentially extending nascent RNA studies to a variety of model systems.

## RESULTS

### fastGRO yields comprehensive nascent transcriptome data in human cells

We obtained fastGRO libraries using the suspension cell line THP-1. These widely used leukemic cells are poorly differentiated myeloid progenitors that can respond to inflammatory stimuli (such as Lipopolysaccharide, LPS) similar to monocytes (Bosshart and Heinzelmann 2016). We processed both unstimulated and LPS-treated THP-1 cells by, first, incubating whole cells in hypotonic solution to cause swelling and subsequent lysis of the plasma membrane (Fig. 1a). Next, we used isolated nuclei to perform *in vitro* run-on reactions with the addition of 4-S-UTP in lieu of the brominated or biotinylated ribonucleotides used in GRO-seq and PRO-seq, respectively. NTPs containing a reactive thiol group are efficiently incorporated by RNA polymerase as evidenced by techniques such as 4SU-seq and TT-seq (Schwanhausser, Busse et al. 2011; Schwalb, Michel et al. 2016) that rely on metabolic labelling starting from thio-nucleoside analogs. In fact, we noticed that RNA yield after affinity purification was in the order of several micrograms, as compared to the yield of GRO-seq and PRO-seq that falls below the most sensitive detection methods. Following isolation by Trizol, we subjected RNA to mild sonication (Fig. 1b-c) using a Bioruptor device. This step is necessary to improve efficiency of the downstream immunopurification and to obtain an even representation of the fully unprocessed mRNA transcript (Supplementary Fig. 1). In fact, we observed that undersonicated RNA resulted in significant loss of resolution at the 3′ of most genes (Supplementary Fig. 1). The incorporated 4-S-UTPs are covalently linked to biotin using a pyridyldithiol-biotin compound. The reaction forms a reversible disulfide bridge between biotin and the uracil base and allows enrichment of bona fide nascent RNA by affinity purification via streptavidin-conjugated beads (Fig. 1a). Lastly, affinity-bound molecules are eluted with harsh reducing conditions to cleave off the biotin adduct and recover nascent RNA fragments that will be incorporated into directional (stranded) Illumina-compatible libraries (see Supplementary Methods for detailed protocol).

**Figure 1.**
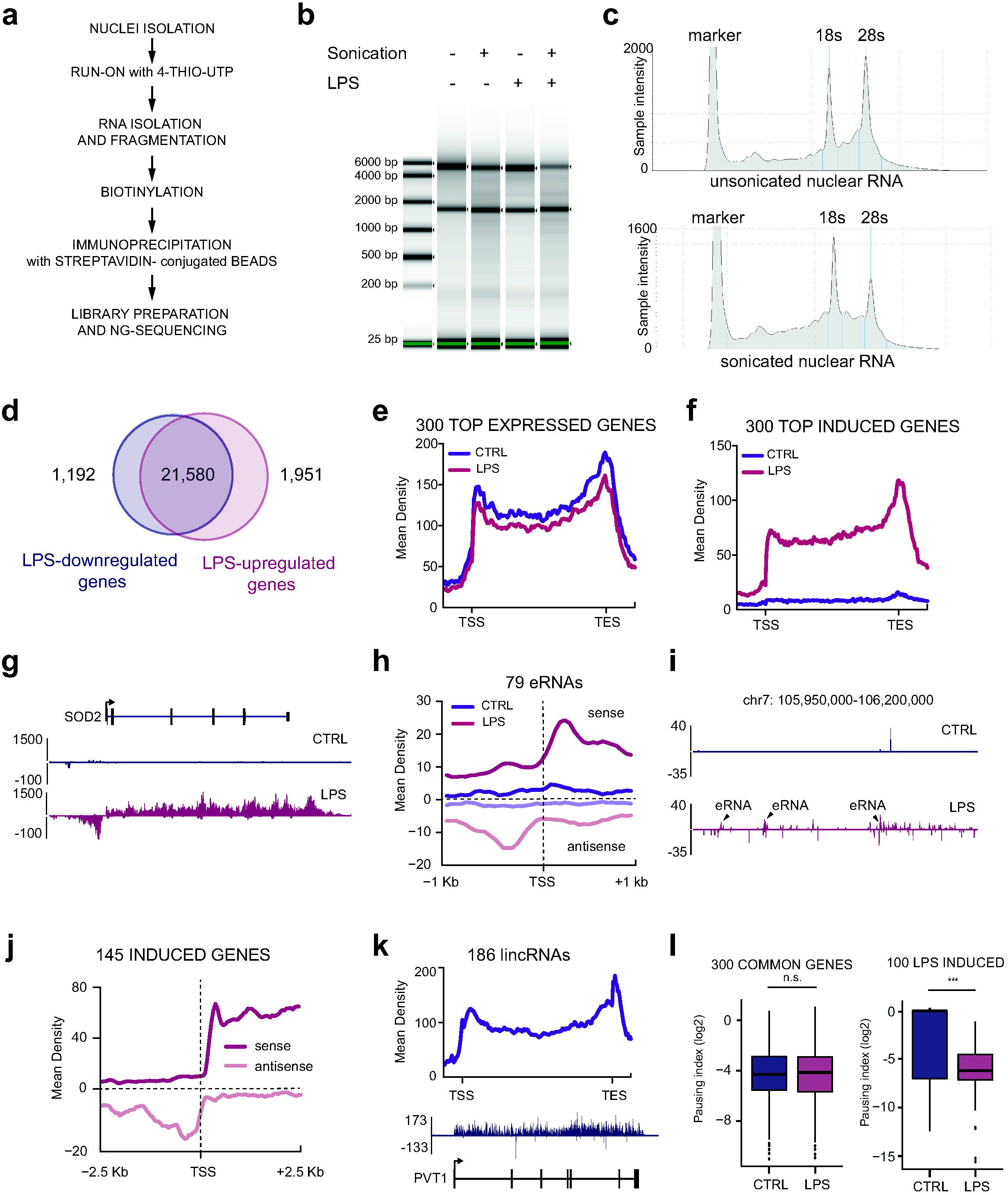
fastGRO generates global nascent transcriptome data. (a) Schematic of fastGRO procedure. Nuclei are first isolated and nuclear run-on (NRO) is performed *in vitro* in the presence of 4-thio-UTP. NRO RNA is isolated and fragmented, biotinylated, recovered using streptavidin-conjugated beads and processed for library preparation. (b-c) Examples of TapeStation run showing mild fragmentation of NRO RNA extracted from LPS-treated and untreated human THP1 cells. (d) CTRL (purple) and LPS-treated THP1 (dark pink) fastGRO samples were analyzed by HOMER to identify common and LPS-induced transcripts. (e) Average density profiles of fastGRO signals for CTRL and LPS-induced THP1 at 300 most expressed genes. (f) Average density profiles of fastGRO reads for CTRL and LPS-induced THP1 at 300 most LPS-induced genes. (g) Screenshot of region surrounding the LPS-induced gene SOD2 showing fastGRO reads along gene body, post-transcription end site and promoter antisense. (h) Average density profiles of sense and antisense fastGRO reads at 79 putative enhancer regions, identified by the level of H3K27ac (Supplemental FigureS3). (i) Screenshots of LPS-induced enhancer RNAs. (j) Average density profile of sense and antisense fastGRO reads at the TSS region of 145 LPS induced genes showing. (k) Average profile of fastGRO reads from untreated THP1 at 186 long intergenic non-coding RNA (lincRNA) and screenshot of the PVT1 lincRNA in untreated THP1 as depicted by fastGRO. (l) Pausing index was calculated from fastGRO reads for 300 highly expressed genes and 300 LPS-induced genes, showing how fastGRO is a useful approach to study RNAPII elongation.

We initially assessed the coverage of fastGRO sequencing by using de novo transcript identification with the HOMER suite (Heinz, Benner et al. 2010). In this analysis, we retrieved nearly 23,000 newly annotated, independent transcripts in both unstimulated and LPS-stimulated THP-1 cells (Fig. 1d). Over 90% of transcripts were expressed at similar levels between both conditions and 1,951 were upregulated by LPS (as compared to 1,192 that were downregulated). Importantly, we assessed that fastGRO is a highly reproducible technique, since replicated experiments that were independently performed show highly significant correlation (Supplementary Fig. 2). Average read profiles at the 300 most expressed genes (Fig. 1e) show a robust signal with seamless coverage along the entire gene body, including the 3′ post-termination region (data were normalized using spike-in of Drosophila RNA, see Methods for details). Furthermore, fastGRO detected strong nascent transcription at LPS-induced genes as well as LPS-related enhancer and super-enhancer sites (Fig. 1f-I, Supplementary Fig. 3). On average, fastGRO proficiently detects nascent RNA at all RNAPII sites, including antisense promoter transcripts that are known to be rapidly degraded by the exosome (Fig. 1j) (Flynn, Almada et al. 2011). Low-expressed lincRNAs were also well represented in our dataset (Fig. 1k). These transcripts are highly regulated and contribute to the expression of neighboring genes, as in the case of PVT1 (Fig. 1k) that is adjacent to the MYC locus and essential for MYC-driven oncogenesis(Tseng, Moriarity et al. 2014). LincRNAs are conventionally defined as noncoding transcripts longer than 200bp. Importantly, even shorter noncoding RNAs, either RNAPII or RNAPIII-dependent, were robustly detected by fastGRO. For instance, we were able to profile tRNAs, snoRNAs and UsnRNAs (Supplementary Fig. 4).

Similar to GRO-seq and PRO-seq, fastGRO profiles are a reflection of RNAPII occupancy and incorporate information on polymerase activity, such as the rate of pause-release and elongation. We used our dataset to calculate pausing indexes (the read ratio between the proximal promoter and the remaining gene body) at 300 highly expressed, constitutive genes and at a group (100) of LPS targets (Fig.1l). Our data show no significant changes (+ or - LPS) in the control group, while the pausing index of LPS-responsive genes decreases dramatically upon stimulation, suggesting steady accumulation of RNAPII into the gene body.

### Unbiased recovery of unprocessed transcripts by fastGRO

Unlike steady-state RNA-seq, nascent RNA-seq captures transcripts before they have been fully processed. Since the vast majority of eukaryotic protein coding genes contain multiple introns, splicing is deemed one of the most frequent and abundant RNA processing events. Therefore, we sought to measure residual splicing events in fastGRO data, to probe the actual enrichment of nascent, unprocessed transcripts. We used MAJIQ (Vaquero-Garcia, Barrera et al. 2016) to determine the relative frequency of splicing junctions, normalized by transcriptome coverage. We stacked up fastGRO of THP1 cells against ribodepleted and poly(A)-selected RNA-seq data that we obtained from the same batch of cells (Fig. 2a). As expected, poly(A) RNA-seq data bear the highest fraction of spliced transcripts (median >80%) as opposed to 20% of fastGRO. Ribo-depleted RNA-seq also carries significantly more junctions, albeit slightly lower than poly(A) RNA-seq (as expected, ribodepletion allows minimal retention of non-polyadenylated, unprocessed RNA species). Next, we compared fastGRO to previously published techniques that capture nascent transcription by means of run-on assay (GRO-seq, PRO-seq), metabolic labeling (TT-seq) and isolation of RNAPII/RNA complexes (NET-seq). By taking advantage of commercially available library preparation kits and a single round of biotin enrichment, fastGRO samples can be prepared within 2.5 days, while most other protocols require 5 days of processing time before obtaining sequencing-ready libraries (Fig. 2b). We mined public repositories for previously published nascent transcriptomic data of THP-1 cells. We retrieved datasets of GRO-seq, PRO-seq and TT-seq. First, we compared the relative enrichment of nascent, unspliced transcript using MAJIQ. Strikingly, fastGRO showed the least contamination of spliced RNA of all techniques (Fig. 2c). Next, we compared the average read density profile of fastGRO, GRO-seq, PRO-seq and TT-seq (normalized by sequencing depth). Across a group of highest expressing protein coding genes, fastGRO and TT-seq similarly displayed smooth and continuous density profiles across the entire gene body (Fig. 2d-e). PRO-seq and GRO-seq profiles appeared more irregular and biased towards the 5′ promoter proximal region (Fig. 2d-e). Since both GRO-seq and PRO-seq protocols comprise multiple size selection steps performed on polyacrylamide gels, they are more likely to introduce a size bias towards smaller RNA fragments.

**Figure 2.**
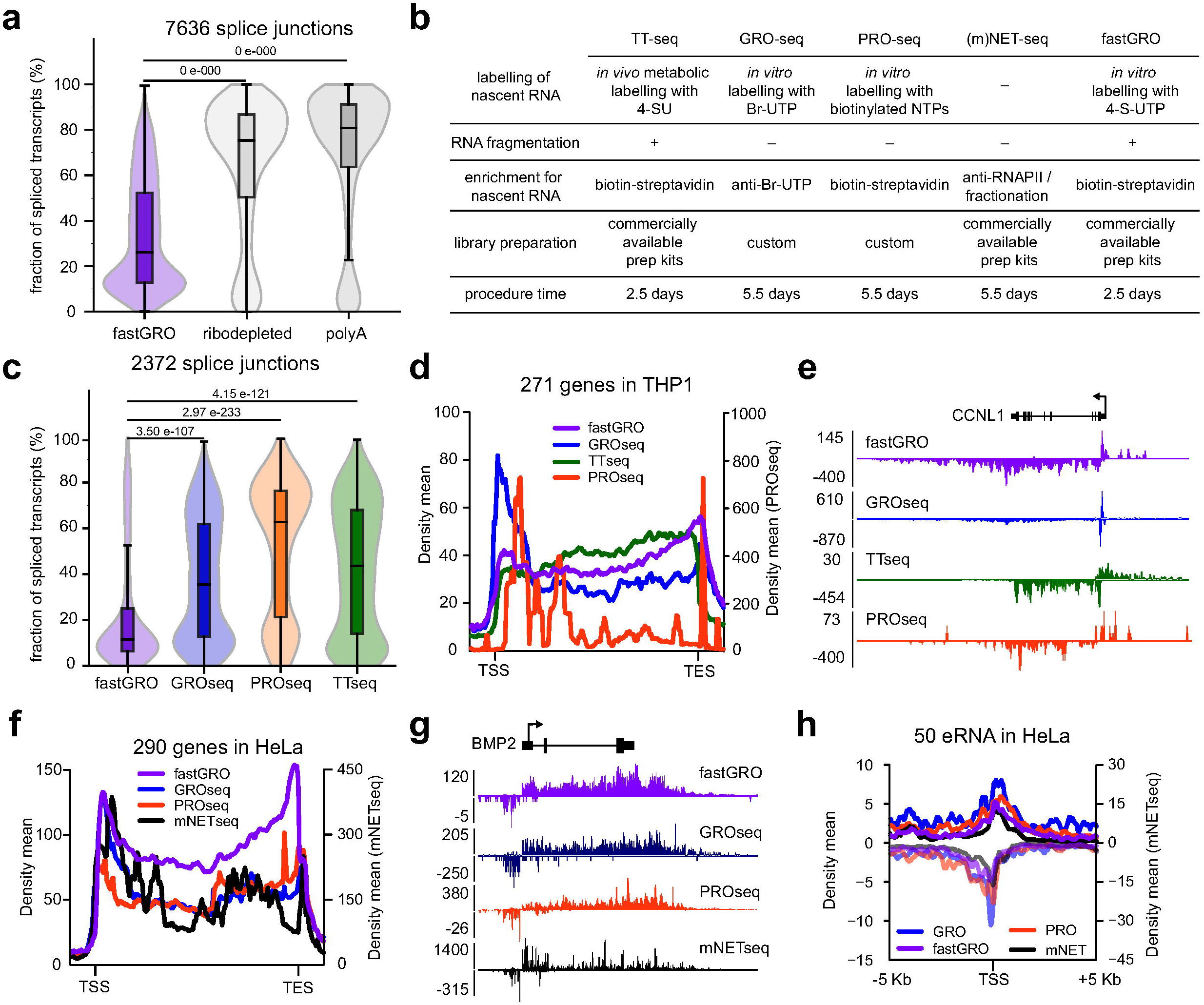
fastGRO recovers nascent, unprocessed and short-lived transcripts. (a) Splice junction analysis by MAJIQ shows substantial recovery of processed RNA by rRNA-depleted (grey) and polyA-enriched RNA-seq (dark grey). fastGRO (purple) is significantly enriched for nascent, unspliced RNA. (b) Comparison of fastGRO to other nascent RNA techniques. An advantage of fastGRO is the overall short processing time (2.5 days, using commercially available library prep kits). (c) fastGRO (purple) shows lower enrichment of spliced junctions than comparable nascent RNA-seq techniques such as GRO-seq (blue), PRO-seq (orange) and TT-seq (green) in THP1 cells. (d) Average profiles of fastGRO, GRO-seq, PRO-seq, TT-seq at 271 highly expressed genes in THP1 cells. fastGRO shows a lower bias toward the 5’ end compared to GRO-seq and recovers more post-termination RNA compared to TT-seq. (e) Screenshot of the CCNL1 gene comparing fastGRO, GRO-seq, PRO-seq and TT-seq in THP1 cells. (f) Average profiles of fastGRO (purple), GRO-seq (blue), PRO-seq (orange), mNET-seq (black) at 290 highly expressed genes in HeLa cells. fastGRO shows a homogenous profile along the whole gene body. (g) Screenshot of the BMP2 gene showing the comparison of fastGRO, GRO-seq, PRO-seq and mNET-seq tracks in HeLa cells. mNET-seq data are downscaled (right y-axis). (h) Average profile of fastGRO, GRO-seq, PRO-seq and mNET-seq reads at 50 eRNAs in HeLa cells. fastGRO recovers bidirectional short-lived eRNAs. mNET-seq data are downscaled (right y-axis).

To ensure that fastGRO is applicable to other cell systems and further the comparison to similar techniques, we generated libraries from HeLa cells. fastGRO showed the most homogeneous coverage across the whole gene body of the top 300 expressed genes (Fig. 2f-g). PRO-seq and GRO-seq profiles showed more 5′ bias and scattered coverage (Fig. 2f-g). We also analyzed an available mNET-seq dataset. The signal was more robust than all other techniques (as per read depth normalization) but heavily scattered due to the nature of NET-seq technology that favors discovery of polymerase pausing sites. Furthermore, fastGRO data ensured comprehensive coverage of bidirectional enhancer RNAs (Fig 2h), comparable to other techniques.

### fastGRO maps the fate of RNA Polymerase post-termination

We observed increased coverage of 3′ regions by fastGRO at several protein coding genes (Fig. 2d-e). In particular, we noticed a robust pile-up of sequencing reads surrounding the annotated transcription end site (TES). Upon recognition of the polyadenylation site (PAS), the Cleavage and Polyadenylation machinery is recruited by RNAPII, resulting in the release of a full-length, capped mRNA precursor that will be handed over to Poly(A) polymerase (Shi and Manley 2015; Proudfoot 2016). However, RNAPII moves further downstream and elongates the post-termination uncapped 3′ RNA, which is promptly degraded by the Xrn2 exonuclease. Running after polymerase for several hundreds of nucleotides, Xrn2 eventually prompts RNAPII arrest and unload from its chromatin template (Eaton and West 2018). Post-termination RNA is rapidly degraded, hence difficult to recover, but provides unique insight into the mechanisms and protein complexes that oversee termination. To fare nascent RNA protocols on their ability to recover 3′ RNA, we generated a ‘termination index’ for all high-expressed genes, by calculating the ratio of normalized read density after and before the annotated PAS (Fig. 3a). We found that fastGRO allowed far more significant recovery of post-termination RNA than PRO-seq and TT-seq (and similar to GRO-seq) (Fig. 3a). This was also evident by plotting read-density profiles centered around the TES of the top expressed transcripts (Fig. 3b) and by looking at specific genes that present exceptionally extended 3′ end such as FUT4 (Fig. 3b-c).

**Figure 3.**
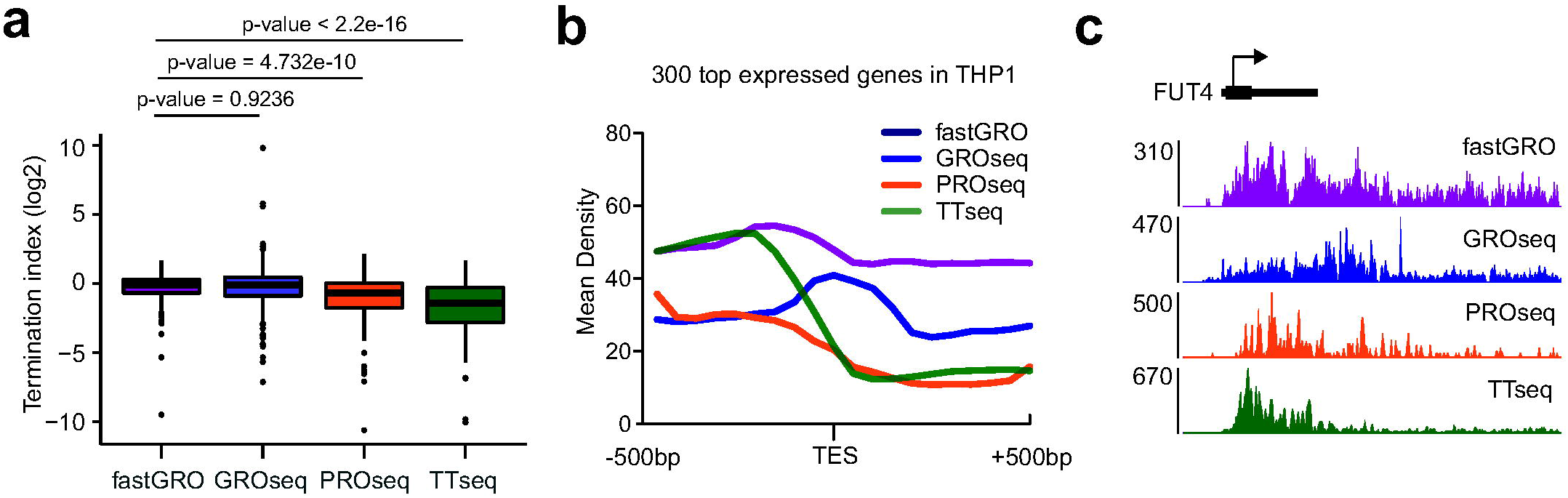
fastGRO identifies transient RNA downstream the poly(A) signal. (a) Box plot of termination index (TI, calculated as ratio between number of reads post Transcription End Site (−1Kb/TES) and number of reads pre TES (TES/+1Kb)) at 300 most expressed gene calculate from fastGRO (purple), GRO-seq (blu), PRO-seq (orange) and TT-seq (green) data. fastGRO and GRO-seq have a comparable TI, while TI generated from PRO-seq and TT-seq are lower (lower coverage of post-termination RNA). (b) Average profile of reads around TES of 300 highly expressed genes calculated for fastGRO, GRO-seq, PRO-seq and TT-seq. fastGRO shows the highest and most homogeneous profile pre- and post-TES compare to other techniques used to study nascent RNA. (c) Screenshot of monocytic gene FUT4 showing the high coverage of post-termination RNA retrieved by fastGRO.

### A scalable global run-on assay

Nascent RNA techniques require a large amount of starting material, restraining their applicability to easily cultured, inexpensive cell types (Wissink, Vihervaara et al. 2019). The recommended starting cell number for optimal GRO-seq and PRO-seq experiments ranges from 1.5 to 2 x 10^7 (Gardini 2017; Wissink, Vihervaara et al. 2019). Similarly, NET-seq is optimized for 1×10^7 cells (Mayer and Churchman 2016), while mNET-seq requires up to 1.6×10^8 starting cells due to the additional RNAPII IP step (Nojima, Gomes et al. 2016). TT-seq, which is based on metabolic labeling, necessitates 300 μ streptavidin immunopurification, equaling up to 3 x 10^7 starting cells (Schwalb, Michel et al. 2016). We initially obtained fastGRO datasets employing 1.5 to 2 x 10^7 cells (Fig. 1-2). In our first attempts to reduce input material, we observed that the use of fewer than 5×10^6 HeLa almost invariably resulted in undetectable amounts of RNA after immunoprecipitation and poor-quality libraries (Supplementary Fig. 5a). We reasoned that boosting the efficiency of thio-UTP biotinylation could improve RNA recovery and the overall quality of sequencing libraries. Therefore, we developed a low-input variant of fastGRO (Supplementary Methods) by taking advantage of the recently optimized methane thiosulfunate biotin (Biotin-MTS) (Duffy, Rutenberg-Schoenberg et al. 2015). While biotin-MTS is more efficient in forming disulfide bonds with 4-S-UTP, it may also cross-react with non thiolated UTP, exposing the entire procedure to contamination of unlabeled, steady state RNA (Marzi, Ghini et al. 2016). In fact, we observed increased carryover of processed RNA when using large number of cells (1.5×10^7). We measured spliced RNA content with MAJIQ and found that biotin-MTS samples carry significantly more contamination than samples prepared with biotin-HPDP (Supplementary Fig. 5b). However, the relative contamination of processed RNA was much reduced in small scale experiments, due to lower RNA concentration in the reaction (Supplementary Fig. 5c). Hence, we adjusted the fastGRO protocol for smaller reaction volumes (we named the low input variant fastGRO-LI) and we generated, first, a scale-down dataset using 5 x 10^6 HeLa cells. Read density profiles of the top 200 genes and the top induced EGF genes showed continuous coverage across the gene body, without loss of resolution as compared to the 2×10^7 cells dataset (Fig. 4a-b). Next, we set up an extended downscale experiment using 2.5, 1 and 0.5x 10^6 cells (8- to 40-fold less than the original experiment). We gauged the fraction of high-input annotated transcripts that were still detected in the low-input samples. Protein coding genes that were undetectable equaled 10% in the 2.5 and 1 x 10^6 cell experiments, and up to 14% in the 0.5. x 10^6 cells sample (Fig. 4c). The vast majority of transcripts were reliably detected by fastGRO-LI and read density profiles were remarkably consistent with those of high-input fastGRO experiments, with loss of resolution at the post-termination RNA (Fig. 4d-e). Taken together, we demonstrate the feasibility of fastGRO with less than a million cells, potentially extending nascent transcriptome analysis to a wider pool of model systems and experimental conditions.

**Figure 4.**
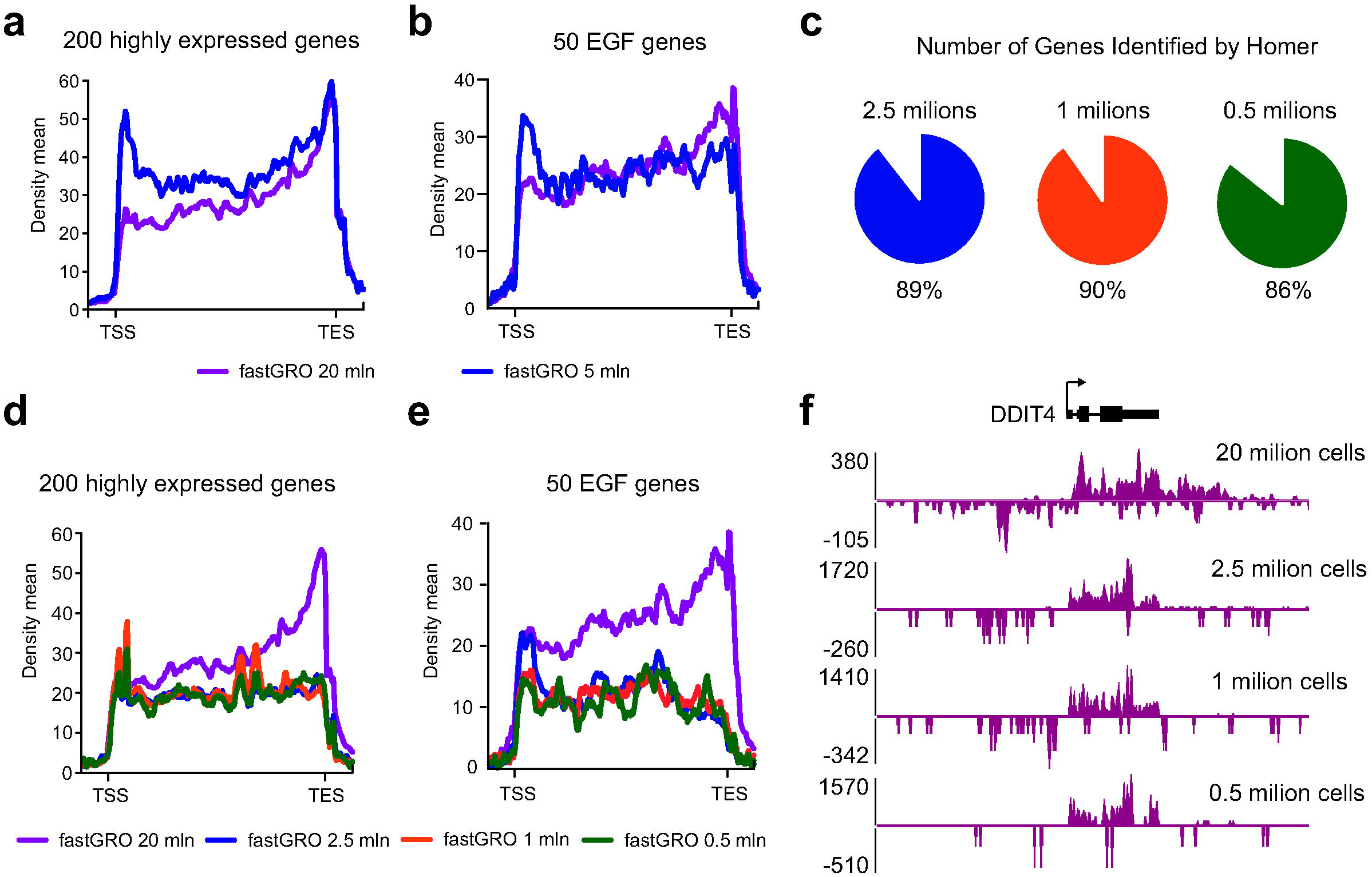
Low-input fastGRO. (a-b) Average profile of fastGRO reads at 200 highly expressed genes (a) and 50 EGF genes (b) in EGF-treated HeLa cells obtained from 20 million (purple) and 5 million cells (blue), showing that fastGRO can be performed with low number of cells using standard protocol and HPDP biotin. (c) Pie charts show the percentage of genes with FPKM>0 in fastGRO obtained with 20 million EGF-induced HeLa cells that have FPKM>0 in fastGRO obtained from 2.5 million (89%), 1 million (90%) and 0.5 million (80%) EGF-induced HeLa cells. (d-e) Average profile of fastGRO reads at 200 highly expressed genes (d) and 50 EGF genes (e) in EGF-treated HeLa cells comparing data obtained from 20 million with HPDP-biotin (purple) and 2.5 million (blue), 1 million (orange) and 0.5 million cells (green) obtained with the fastGRO-LI protocol and MTS-biotin. (f) Screenshot of DDIT4 gene showing fastGRO tracks obtained using either standard (20 million cells) and fastGRO-LI protocols (2.5, 1, and 0.5 million cells).

## Discussion

We developed a fast protocol to generate genomic libraries of nascent RNA, based on a nuclear run-on assay. During the run-on reaction, newly synthesized RNA incorporates the ribonucleotide analog 4-S-UTP (Fig. 5). Sulfhydril-reactive biotin is then covalently bound to UTP analogs, allowing the affinity-based purification of fragmented nascent transcripts. After elution with a harsh reducing buffer, RNA is subjected to directional library preparation as per Illumina guidelines (Fig. 5). fastGRO generates comprehensive and seamless coverage of RNAPII (and RNAPIII) activity for the most abundant RNA species (highly expressed protein coding genes, small noncoding RNAs) as well as those harder to detect (antisense promoter transcripts, eRNAs, lincRNAs).

**Figure 5.**
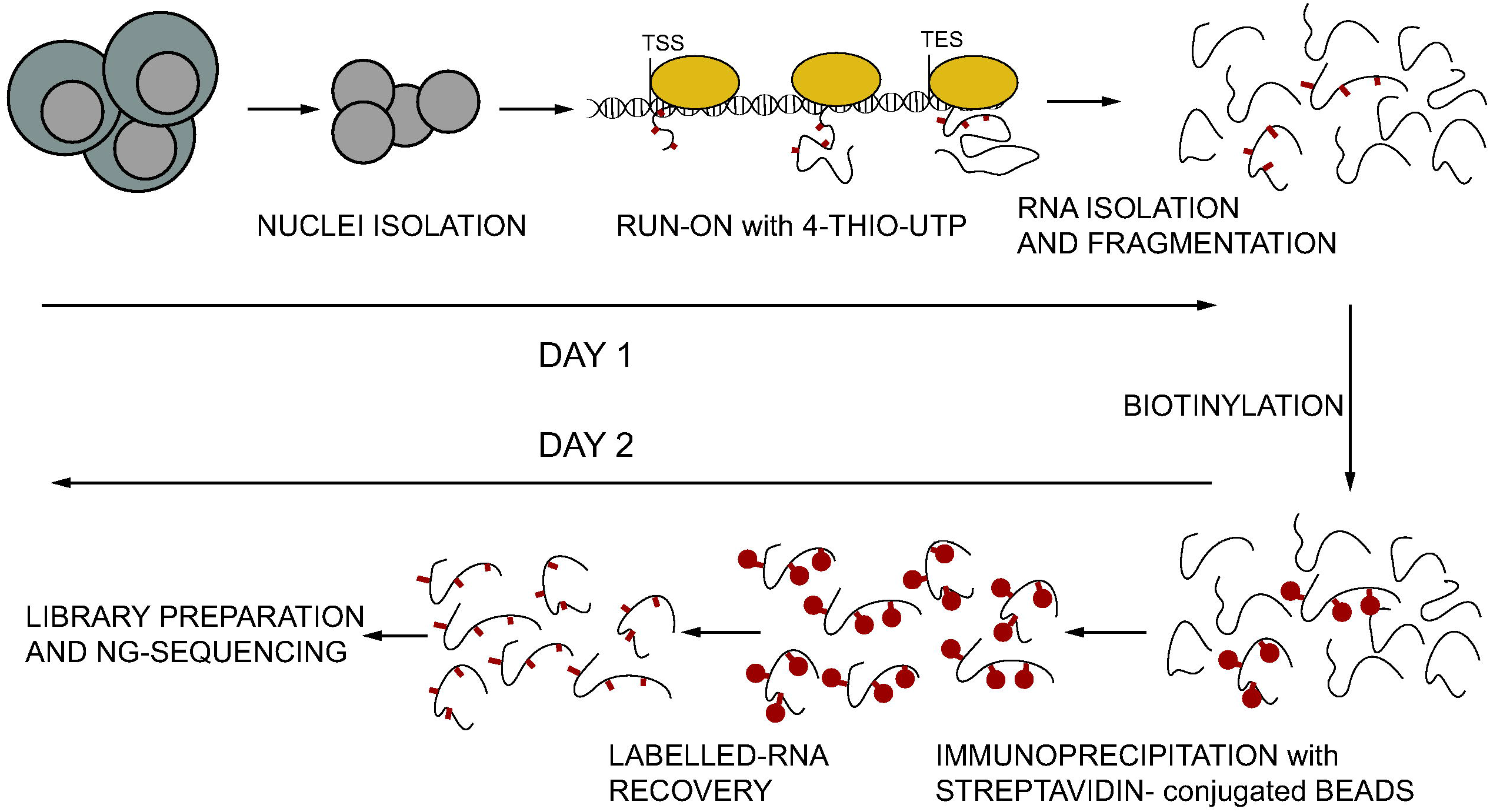
Overview of fastGRO. In day 1, nuclei are isolated and *in vitro* run-on is performed in a solution containing 4-thio-UTP that is incorporated in nascent RNA. After isolation using Trizol and ethanol precipitation, RNA is fragmented and snap-frozen. In day 2, 4-thio-UTP containing RNA is biotinylated using either HPDP- (standard protocol) or MTS-biotin (fastGRO-LI protocol for low input sample) and recovered by immunoprecipitation using streptavidin-conjugated beads. Labeled RNA is recovered by elution in DTT solution, purified and used for NGS libraries preparation with commercially available kits.

We show that fastGRO offers practical advantages over similar techniques, such as GRO-seq and PRO-seq, that are frequently employed to gauge the dynamics and processivity of RNAPII: I) the overall sample processing time is reduced by more than half; II) custom library preparation is replaced by Illumina-compatible prep kits that are streamlined and generate more reproducible libraries (cutting off the user-dependent size selection steps). Furthermore, fastGRO yields significantly better enrichment of nascent, genuinely unprocessed RNA. In fact, contamination of spliced RNA in fastGRO is the lowest among all nascent RNA-seq types, including run-on based techniques as well as metabolic labelling methods (i.e. TT-seq). A superior enrichment of unprocessed, short-lived transcripts is also seen at the post-termination site of protein coding genes. After encountering the first polyadenylation signal, RNAPII continues its course for hundreds to thousands of kilobases, until Xrn2-dependent termination effectively dislodges the enzyme off of its DNA template. We calculated a termination index for a set of highly expressed genes, to demonstrate that fastGRO provides better coverage of RNAPII activity past the polyadenylation site.

Lastly, we established that fastGRO can be performed with lower amounts of input material. A major limitation of all current methods of nascent RNA sequencing is the requirement of 10 to 30 million cells per single library. We performed a serial downscale of fastGRO using a modified protocol (fastGRO-LI) that employs a highly reactive biotin conjugate (biotin-MTS). Our results suggest that fastGRO can be performed with as few as 0.5 x 10^6 cells (30 to 60 times fewer than GRO-seq, PRO-seq or TT-seq), at the expense of less than 15% of the transcriptome. This may bring about, for the first time, portability of nascent RNA technology to a wider range of model systems, including primary human and mouse cells.

Nascent RNA-seq is a method of choice when dissecting RNAPII regulatory steps (such as pausing, elongation and termination). Additionally, it provides the most accurate quantitative data on gene regulation, since it reflects the real-time activity of polymerase. In addition to protein coding genes, nascent RNA-seq methods are also employed to measure RNA species that are undetectable or poorly represented in canonical RNA-seq datasets. There has been a recent research focus on unstable and/or low-abundant RNA categories, which has propelled RNA biology and impressed a new mark on the fields of epigenetics and transcription regulation. For instance, the low-expressed and poorly evolutionarily-constrained lincRNAs were found to regulate a variety of biological processes by associating with either repressing or activating chromatin complexes (Gardini and Shiekhattar 2015; Ransohoff, Wei et al. 2018). Additionally, eRNAs have been used to gauge enhancer activity throughout development and were shown to directly impact gene expression (Andersson, Gebhard et al. 2014; Lai, Gardini et al. 2015; Bose, Donahue et al. 2017). Lastly, antisense promoter transcripts are rapidly degraded but offer an invaluable insight into the architecture of eukaryotic promoters and the mechanisms of productive elongation and early termination (Almada, Wu et al. 2013; Andersson, Chen et al. 2015; Jin, Eser et al. 2017). Nascent RNA methods provide existential support to these lines of research and fastGRO represents a standardized, user-friendly and scalable technique that can be integrated into several experimental settings.

## Material and Methods

### Cell lines

THP-1 were obtained from American Type Culture Collection (ATCC) and maintained in Roswell Park Memorial Institute (RPMI)-1640 medium (Corning) supplemented with 10% (v/v) of super calf serum (GEMcell) and 2 mM of L-glutamine (Corning). HeLa were obtained from ATCC and maintained in Dulbecco’s Modified Eagle’s Medium (DMEM) supplemented with 10% super calf serum (GEMcell) and 2 mM L-glutamine (Corning). THP1 cells were treated with 5 µg/ml of LPS (Invitrogen) for 4 hours in growing medium. HeLa were treated with 100 ng/ml of rEGF (Invitrogen) for 15 minutes in growing medium.

### fastGRO

20-5 million of cells were washed twice with ice-cold PBS before adding swelling buffer (10 mM Tris-HCL pH 7.5, 2mM MgCl_2_, 3 mM CaCl_2_, 2U/ml Superase-in (Invitrogen)). Cells were swelled for 5 min on ice, washed with swelling buffer + 10% glycerol and then lysed in lysis buffer (10 mM Tris-HCL pH 7.5, 2 mM MgCl_2_, 3 mM CaCl_2_, 10% glycerol, 1 ml Igepal (NP-40), 2 U/ml Superase-in) to isolate nuclei. Nuclei were washed twice with lysis buffer and resuspended in freezing buffer (40% glycerol, 5 mM MgCl_2_, 0.1 mM 0.5M EDTA, 50 mM Tris-HCL pH 8.3) to a concentration of 2×10^7 nuclei per 100 µL. Nuclei were then frozen in dry ice and stored at −80 °C. Nuclei were thawed on ice and an equal volume of pre-warmed nuclear run-on reaction buffer (10 mM Tris-HCl pH 8, 5 mM MgCl_2_, 300 mM KCl, 1 mM DTT, 500 µM ATP, 500 µM GTP, 500 µM 4-thio-UTP, 2 µM CTP, 200 U/ml Superase-in, 1% Sarkosyl (N-Laurylsarcosine sodium salt solution) was added and incubated for 7 min at 30 °C for nuclear run-on. Nuclear run-on RNA was extracted with TRIzol LS reagent (Invitrogen) following the manufacturer’s instructions and ethanol precipitated. NRO-RNA was resuspended in water and concentration was determined with Qubit High Sensitivity Assay kit (Invitrogen). Up to 150 μg of RNA was transfer to a new tube and 5-10% of spike-in RNA was added. RNA was then fragmented with a Bioruptor UCD-200 for 1-5 cycles of 30 seconds ON / 30 seconds OFF, high settings. Fragmentation efficiency was analyzed by running fragmented and unfragmented RNA on Agilent 2200 TapeStation using High Sensitivity RNA ScreenTapes following manufacturer’s instructions. Fragmented RNA was incubated in Biotinylation Solution (100 mM Tris pH 7.5, 10 mM EDTA pH 8.0, 40% dimethylformamide, 200 μg/ml EZ-link HPDP Biotin (Thermo Scientific)) for 2h in the dark at 25 °C, 800 rpm. After ethanol precipitation, the biotinylated RNA was resuspended in water and biotinylated-RNA was separated with M280 Streptavidin Dynabeads (Invitrogen). 100 ul/sample of beads were washed twice with 2 volumes of freshly prepared wash buffer (100 mM Tris pH 7.5, 10 mM EDTA pH 8.0, 1M NaCl, 0.1% (v/v) Tween-20) and resuspended in 1 volume of wash buffer and added to the biotinylated-RNA. After 15 minutes in rotation at 4 °C, beads were washed three times with wash buffer pre-warmed at 65 °C and three times with room temperature wash buffer. 4-SUTP containing RNA was eluted in 100 mM DTT buffer and purified with RNA Clean and Purification kit (Zymo Research) with in-column DNAseq reaction to eliminate traces of genomic DNA. The eluted RNA was quantified with Qubit High Sensitivity Assay kit (Invitrogen) and used to produce barcoded RNA sequencing libraries using the NEBNext Ultra II Directional RNA Library Prep kit (New England Biolabs). Libraries were sequenced on Illumina NextSeq 500.

### Low-input fastGRO (fastGRO-LI)

5-0.5 million nuclei were extracted as described for fastGRO and resuspended in freezing buffer (40% glycerol, 5 mM MgCl_2_, 0.1 mM 0.5M EDTA, 50 mM Tris-HCL pH 8.3) to a concentration of up to 2×10^6 nuclei per 10 μL. Nuclei were then frozen in dry ice and stored at −80 °C. Run-on reaction was performed as described for fastGRO, NRO-RNA was resuspended in water and concentration was determined with Qubit High Sensitivity Assay kit (Invitrogen). Up to 30 μg of RNA was transfer to a new tube and 5-10% of spike-in RNA was added. RNA was then fragmented with a Bioruptor UCD-200 for 30 seconds, low settings. Fragmentation efficiency was analyzed by running fragmented and unfragmented RNA on Agilent 2200 TapeStation using High Sensitivity RNA ScreenTapes following manufacturer’s instructions. Fragmented RNA was incubated in low-input Biotinylation Solution (25 mM Hepes pH 7.5, 1 mM EDTA, 25% dimethylformamide, 16.4 μM MTS-Biotin (Biotium)) for 30 minutes in the dark at 25 °C, 800 rpm. After ethanol precipitation, the biotinylated RNA was resuspended in water and DNAse treatment was performed with TURBO DNAse (Invitrogen) following manufacturer instructions. Biotinylated-RNA was separated with M280 Streptavidin Dynabeads (Invitrogen): 25 μl/sample of beads were washed twice with 2 volumes of freshly prepared wash buffer (100 mM Tris pH 7.5, 10 mM EDTA pH 8.0, 1M NaCl, 0.1% (v/v) Tween-20) and resuspended in 1 volume of wash buffer and added to the biotinylated-RNA. After 15 minutes in rotation at 4 °C, beads were washed three times with wash buffer pre-warmed at 65 °C and three times with room temperature wash buffer. thio-UTP containing RNA was eluted in 100 mM DTT buffer, ethanol purified used to produce barcoded RNA sequencing libraries using the NEBNext Ultra II Directional RNA Library Prep kit (New England Biolabs). Libraries were sequenced on Illumina NextSeq 500.

### Spike-in preparation

Drosophila S9 cells were incubated for 5 minutes with 50mM of 4-thiouridine (4sU) at room temperature. Cells were then washed twice with 1X PBS, lyzed in Trizol reagent. RNA was extracted with Direct-zol Mini prep kit (Zymo research). Aliquots of 2 µg were prepared, snap-frozen in liquid nitrogen and store at −80 °C.

### Analysis of RNA-seq data

Reads were aligned to hg19 using STAR v2.5 (Dobin, Davis et al. 2013), in 2-pass mode with the following parameters: --quantMode TranscriptomeSAM --outFilterMultimapNmax 10 -- outFilterMismatchNmax 10 --outFilterMismatchNoverLmax 0.3 --alignIntronMin 21 -- alignIntronMax 0 --alignMatesGapMax 0 --alignSJoverhangMin 5 --runThreadN 12 -- twopassMode Basic --twopass1readsN 60000000 --sjdbOverhang 100. The latest annotations obtained from Ensembl were used to build reference indexes for the STAR alignment. Bam files were filtered based on alignment quality (q = 10) using Samtools v0.1.19 (Li, Handsaker et al. 2009). Bam files were then normalized based on the number of reads of spike-in/total read number with Samtools and bigwig files were built with deeptool 3.3.1 (Ramirez, Ryan et al. 2016).

For nascent RNA analysis, bam files were transformed in bed file with bedtools (bamtobed option) and subjected to analysis with HOMER v4.11 (Heinz, Benner et al. 2010). For the identification of new transcripts, findPeaks.pl was used to analyze fastGRO data with the following parameters: -style groseq -tssFold 6 -bodyFold 5 -pseudoCount 0.5 -minBodySize 500 -maxBodySize 100000. To analyse gene expression, FPKM (Fragments Per Kilobase of exon per Million fragments mapped) was calculated with HOMER using analyzeRepeats.pl (parameters: rna -count genes -strand - -rpkm -condenseGenes) and addGeneAnnotation.pl. FPKM were used to analyze differential gene expression levels, normalized by feature length with DESeq2 (Love, Huber et al. 2014).

### PolyA RNA-seq and ribodepleted RNA-seq

Total RNA was extracted using Direct-zol RNA Miniprep kit (Zymo Research). For polyA RNA-seq, the polyA fraction was isolated by running RNA samples through the Oligo(dT) Dynabeads (Invitrogen). For ribodepleted RNA-seq, ribosomal RNA was removed by the KAPA RNA HyperPrep Kit (Illumina). The resulting RNA was subjected to strand-specific library preparation using the SENSE mRNA-Seq Library Prep Kit V2 (Lexogen). Sequencing was performed on Nextseq500 (Illumina).

### Published genome-wide data and analysis

Original high-throughput sequencing data are deposited at the Gene Expression Omnibus with accession number GSE143844. GRO-seq, PRO-seq and TT-seq datasets from THP1 cells can be retrieved under the following accession numbers: GSM2428733, GSM2544240, GSM3681467. GSM3681459 and GSM3681461 datasets were used for H3K27ac ChIP-seq in THP1. GRO-seq, PRO-seq, mNET-seq datasets from HeLa cells can be retrieved under the following accession numbers: GSM2428725, GSM2692352, GSM2357382. Data were downloaded and re-analyzed as described for nascent RNA for GRO-seq, PRO-seq, TT-seq and mNET-seq. H3K27ac ChIPseq data were aligned to hg19, using Burrows Wheeler Alignment tool (BWA), with the MEM algorithm. Aligned reads were filtered based on mapping quality (MAPQ > 10) to restrict our analysis to higher quality and likely uniquely mapped reads, and PCR duplicates were removed.

### Average density analysis, pausing index and termination index analysis

fastGRO, GRO-seq, PRO-seq, TT-seq, mNET-seq and RNA-seq data were subjected to read density analysis after spike-in (for fastGRO) and sequencing depth normalization. seqMINER 1.3.3 was used to extract read densities, and mean density profiles were then generated in R 3.5.3 using ggplot2 (Villanueva and Chen 2019). For pausing index analysis, the ratio between read counts at the TSS (−50/+150 bp) and read counts across the rest of the gene body (+150/termination end site) was calculated. For termination index analysis, the ratio between read counts post the termination end site (TES, TES / +1 kb) and read counts pre TES (−1 kb / TES) were calculated. Statistical robustness was calculated with Wilcoxon rank-sum tests.

### Splice junction analysis

Reads were trimmed to the average length of the reads in the dataset with the shortest reads in each given comparison. Only one of two reads from paired-end data was used in cases involving comparisons between single- and paired-end datasets. Reads were aligned to human genome assembly GRCh38 using STAR (version 2.5.4b)(Dobin, Davis et al. 2013). Splice junctions were identified and quantified using MAJIQ (Vaquero-Garcia, Barrera et al. 2016) requiring a minimum number of reads on average in intronic sites (--min-intronic-cov) of 0.005 and the number of intronic bins with some coverage (--irnbins) of 0.1. Only junctions common to all samples in any given comparison were used in the analyses. Statistical tests performed were Wilcoxon rank-sum tests.

## Acknowledgments

This work was supported by grants from the National Institute of Health (R01 CA202919 to A.G.), the American Cancer Society (RSG-18-157-01 to A.G.) and the G. Harold and Leila Y. Mathers Foundation (MF-1809-00172 to A.G.). All genome-wide data were generated and processed at the Wistar Institute with the outstanding technical support from all members of the Genomics Facility, particularly Dr. Shashi Bala (NIH-NCI funded CCSG: P30-CA010815).

## Supplementary figure legends

**Supplementary Figure S1.** (a) Tapestation profiles of unsonicated and sonicated RNA. Sonication was performed for 30 (short), 60 (medium) and 150 seconds (high). Profiles show decrease in the level of rRNAs (especially 28s rRNA) upon sonication. (b-c) Average profiles of fastGRO reads along the gene body (b) and surrounding the TES (c) of 290 highly expressed genes in HeLa, obtained from samples with different levels of RNA sonication, showing a decrease in coverage at the 3’ region of expressed genes in samples with low sonication (short) compared to more sonicated samples.

**Supplementary Figure S2.** Correlation plot of FPKM for all expressed genes in two replicates of fastGRO in untreated (a) and LPS-induced (b) THP1 cells. The high level of correlation (rho > 0.83) shows high reproducibility of fastGRO.

**Supplementary Figure S3.** Average profile of H3K27ac ChIP-seq data at 79 eRNA regions in untreated THP1 cells. Related to Figure 1h.

**Supplementary Figure S4.** Average profile of fastGRO reads at 127 tRNA (a), 60 snoRNA (b) and 10 UsnRNA (c) expressed in THP1 cells. (d) Screenshot of a region surrounding the BORCS6 gene, showing examples of tRNA (tRNA_Ile) and snoRNA (SNORD118) identified by fastGRO in untreated THP1 cells.

**Supplementary Figure S5.** (a) Screenshot of DDIT4 region showing fastGRO tracks obtained from 20 million, 5 million and 2.5 million cells obtained with the standard protocol using HPDP-biotin, showing that fastGRO in low-input samples cannot be performed using HPDP-biotin. (b) Box plot showing percentage of splice junctions calculated by MAJIQ in HPDP and MTS fastGRO samples from LPS-induced THP1 cells. fastGRO with HPDP-biotin recovers more unprocessed, immature RNA compared to MTS-biotin based fastGRO, as suggested by the lower percentage of splice junctions identified by MAJIQ in HPDP-biotin fastGRO sample. (c) Box plot showing percentage of splice junctions calculated by MAJIQ in MTS fastGRO samples obtained from decreasing number of LPS-induced THP1 cells. The analysis shows that fastGRO performed with MTS-biotin retrieves similar levels of processed RNA independently of the starting number of cells.

## Reference

Adelman, K. and J. T. Lis (2012). “Promoter-proximal pausing of RNA polymerase II: emerging roles in metazoans.” Nat Rev Genet 13(10): 720–731.

Almada, A. E., X. Wu, et al. (2013). “Promoter directionality is controlled by U1 snRNP and polyadenylation signals.” Nature 499(7458): 360–363.

Andersson, R., Y. Chen, et al. (2015). “Human Gene Promoters Are Intrinsically Bidirectional.” Mol Cell 60(3): 346–347.

Andersson, R., C. Gebhard, et al. (2014). “An atlas of active enhancers across human cell types and tissues.” Nature 507(7493): 455–461.

Bose, D. A., G. Donahue, et al. (2017). “RNA Binding to CBP Stimulates Histone Acetylation and Transcription.” Cell 168(1-2): 135–149 e122.

Bosshart, H. and M. Heinzelmann (2016). “THP-1 cells as a model for human monocytes.” Ann Transl Med 4(21): 438.

Core, L. J., A. L. Martins, et al. (2014). “Analysis of nascent RNA identifies a unified architecture of initiation regions at mammalian promoters and enhancers.” Nat Genet 46(12): 1311–1320.

Core, L. J., J. J. Waterfall, et al. (2008). “Nascent RNA sequencing reveals widespread pausing and divergent initiation at human promoters.” Science 322(5909): 1845–1848.

Dobin, A., C. A. Davis, et al. (2013). “STAR: ultrafast universal RNA-seq aligner.” Bioinformatics 29(1): 15–21.

Duffy, E. E., M. Rutenberg-Schoenberg, et al. (2015). “Tracking Distinct RNA Populations Using Efficient and Reversible Covalent Chemistry.” Mol Cell 59(5): 858–866.

Eaton, J. D. and S. West (2018). “An end in sight? Xrn2 and transcriptional termination by RNA polymerase II.” Transcription 9(5): 321-326.

Flynn, R. A., A. E. Almada, et al. (2011). “Antisense RNA polymerase II divergent transcripts are P-TEFb dependent and substrates for the RNA exosome.” Proc Natl Acad Sci U S A 108(26): 10460–10465.

Gardini, A. (2017). “Global Run-On Sequencing (GRO-Seq).” Methods Mol Biol 1468: 111–120.

Gardini, A. and R. Shiekhattar (2015). “The many faces of long noncoding RNAs.” FEBS J 282(9): 1647–1657.

Gariglio, P., M. Bellard, et al. (1981). “Clustering of RNA polymerase B molecules in the 5’ moiety of the adult beta-globin gene of hen erythrocytes.” Nucleic Acids Res 9(11): 2589–2598.

Gariglio, P., J. Buss, et al. (1974). “Sarkosyl activation of RNA polymerase activity in mitotic mouse cells.” FEBS Lett 44(3): 330–333.

Heinz, S., C. Benner, et al. (2010). “Simple combinations of lineage-determining transcription factors prime cis-regulatory elements required for macrophage and B cell identities.” Mol Cell 38(4): 576–589.

Jin, Y., U. Eser, et al. (2017). “The Ground State and Evolution of Promoter Region Directionality.” Cell 170(5): 889–898 e810.

Kwak, H. and J. T. Lis (2013). “Control of transcriptional elongation.” Annu Rev Genet 47: 483–508.

Lai, F., A. Gardini, et al. (2015). “Integrator mediates the biogenesis of enhancer RNAs.” Nature 525(7569): 399–403.

Lai, F. and R. Shiekhattar (2014). “Enhancer RNAs: the new molecules of transcription.” Curr Opin Genet Dev 25C: 38-42.

Li, H., B. Handsaker, et al. (2009). “The Sequence Alignment/Map format and SAMtools.” Bioinformatics 25(16): 2078–2079.

Love, M. I., W. Huber, et al. (2014). “Moderated estimation of fold change and dispersion for RNA-seq data with DESeq2.” Genome Biol 15(12): 550.

Mahat, D. B., H. Kwak, et al. (2016). “Base-pair-resolution genome-wide mapping of active RNA polymerases using precision nuclear run-on (PRO-seq).” Nat Protoc 11(8): 1455–1476.

Marzi, M. J., F. Ghini, et al. (2016). “Degradation dynamics of microRNAs revealed by a novel pulse-chase approach.” Genome Res 26(4): 554–565.

Mayer, A. and L. S. Churchman (2016). “Genome-wide profiling of RNA polymerase transcription at nucleotide resolution in human cells with native elongating transcript sequencing.” Nat Protoc 11(4): 813–833.

Mayer, A., J. di Iulio, et al. (2015). “Native elongating transcript sequencing reveals human transcriptional activity at nucleotide resolution.” Cell 161(3): 541–554.

Nojima, T., T. Gomes, et al. (2016). “Mammalian NET-seq analysis defines nascent RNA profiles and associated RNA processing genome-wide.” Nat Protoc 11(3): 413–428.

Pai, A. A. and F. Luca (2019). “Environmental influences on RNA processing: Biochemical, molecular and genetic regulators of cellular response.” Wiley Interdiscip Rev RNA 10(1): e1503.

Proudfoot, N. J. (2016). “Transcriptional termination in mammals: Stopping the RNA polymerase II juggernaut.” Science 352(6291): aad9926.

Ramirez, F., D. P. Ryan, et al. (2016). “deepTools2: a next generation web server for deep-sequencing data analysis.” Nucleic Acids Res 44(W1): W160–165.

Ransohoff, J. D., Y. Wei, et al. (2018). “The functions and unique features of long intergenic non-coding RNA.” Nat Rev Mol Cell Biol 19(3): 143–157.

Schoenberg, D. R. and L. E. Maquat (2012). “Regulation of cytoplasmic mRNA decay.” Nat Rev Genet 13(4): 246–259.

Schwalb, B., M. Michel, et al. (2016). “TT-seq maps the human transient transcriptome.” Science 352(6290): 1225–1228.

Schwanhausser, B., D. Busse, et al. (2011). “Global quantification of mammalian gene expression control.” Nature 473(7347): 337–342.

Shi, Y. and J. L. Manley (2015). “The end of the message: multiple protein-RNA interactions define the mRNA polyadenylation site.” Genes Dev 29(9): 889–897.

Stark, R., M. Grzelak, et al. (2019). “RNA sequencing: the teenage years.” Nat Rev Genet 20(11): 631–656.

Tseng, Y. Y., B. S. Moriarity, et al. (2014). “PVT1 dependence in cancer with MYC copy-number increase.” Nature 512(7512): 82–86.

Vaquero-Garcia, J., A. Barrera, et al. (2016). “A new view of transcriptome complexity and regulation through the lens of local splicing variations.” Elife 5: e11752.

Villanueva, R. A. M. and Z. J. Chen (2019). “ggplot2: Elegant Graphics for Data Analysis, 2nd edition.” Measurement-Interdisciplinary Research and Perspectives 17(3): 160–167.

Wissink, E. M., A. Vihervaara, et al. (2019). “Nascent RNA analyses: tracking transcription and its regulation.” Nat Rev Genet 20(12): 705–723.

Yamada, T. and N. Akimitsu (2019). “Contributions of regulated transcription and mRNA decay to the dynamics of gene expression.” Wiley Interdiscip Rev RNA 10(1): e1508.

