## Supplementary Figures for "Rapid and scalable profiling of nascent RNA with fastGRO"

SUPPLEMENTARY FIGURE S1

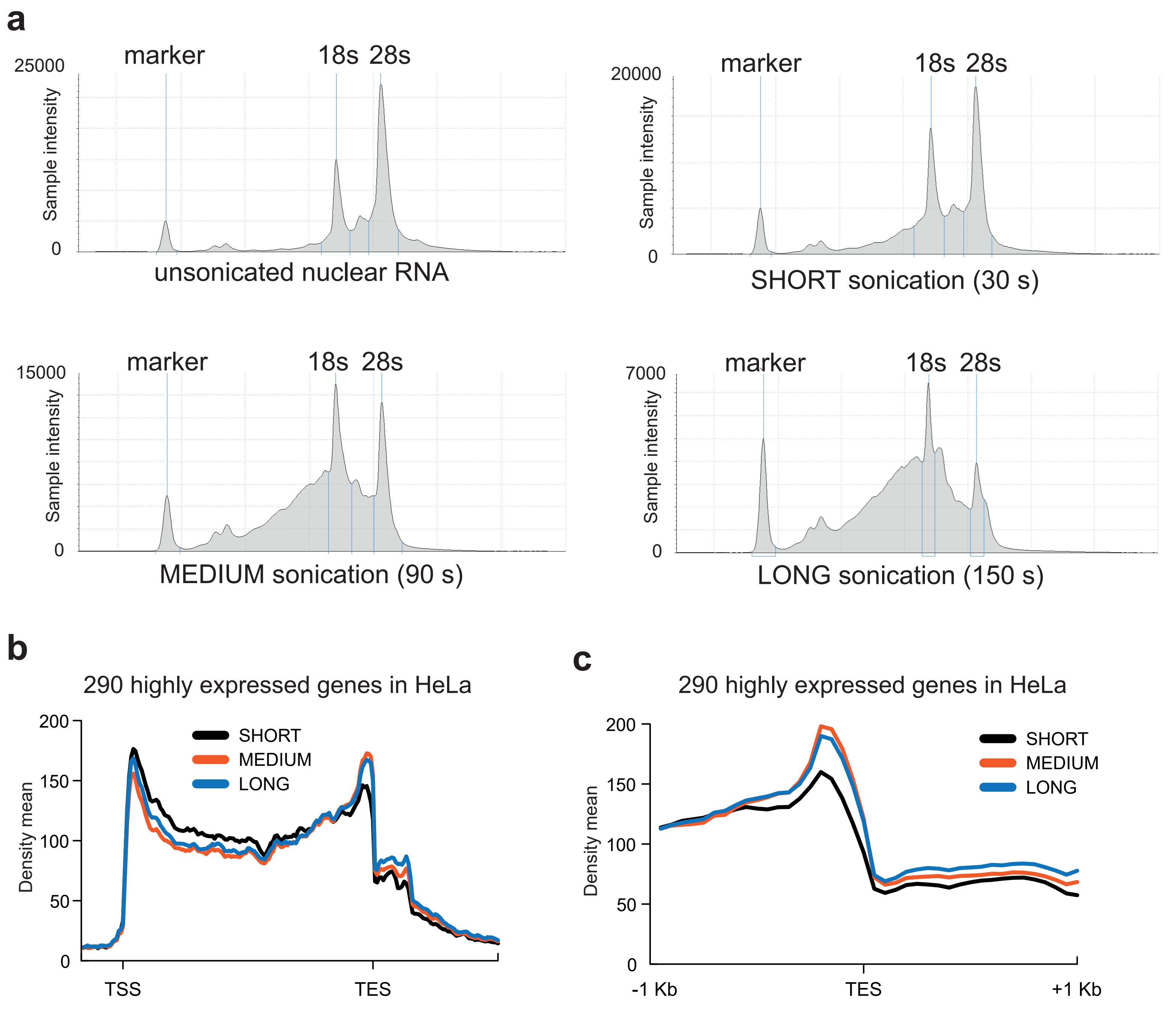

SUPPLEMENTARY FIGURE S2

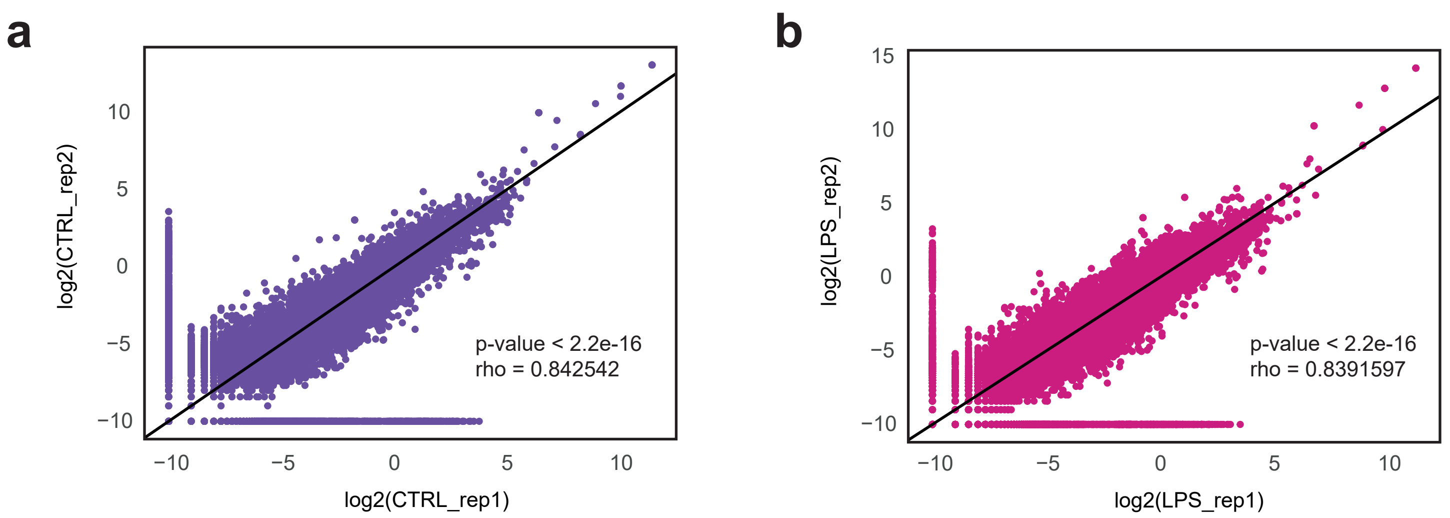

### SUPPLEMENTARY FIGURE S3

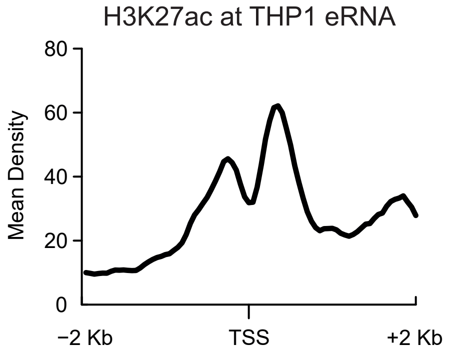

### SUPPLEMENTARY FIGURE S4

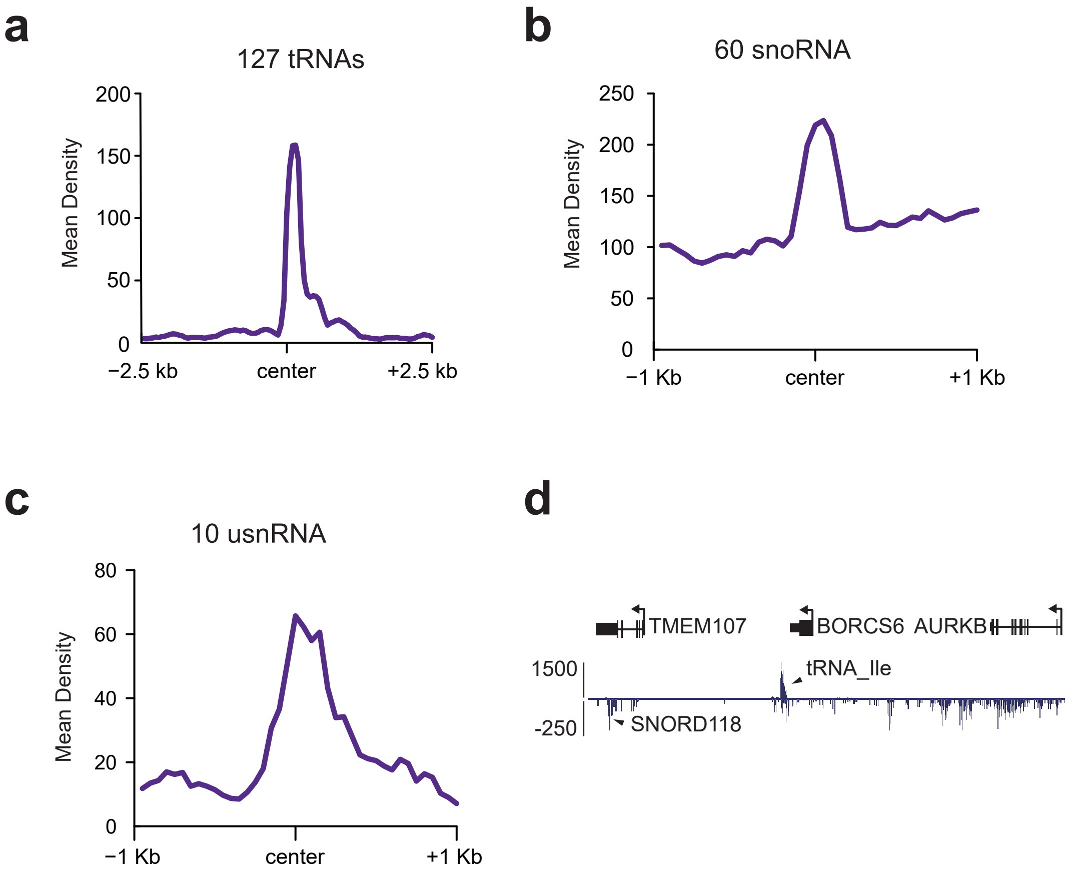

### SUPPLEMENTARY FIGURE S5

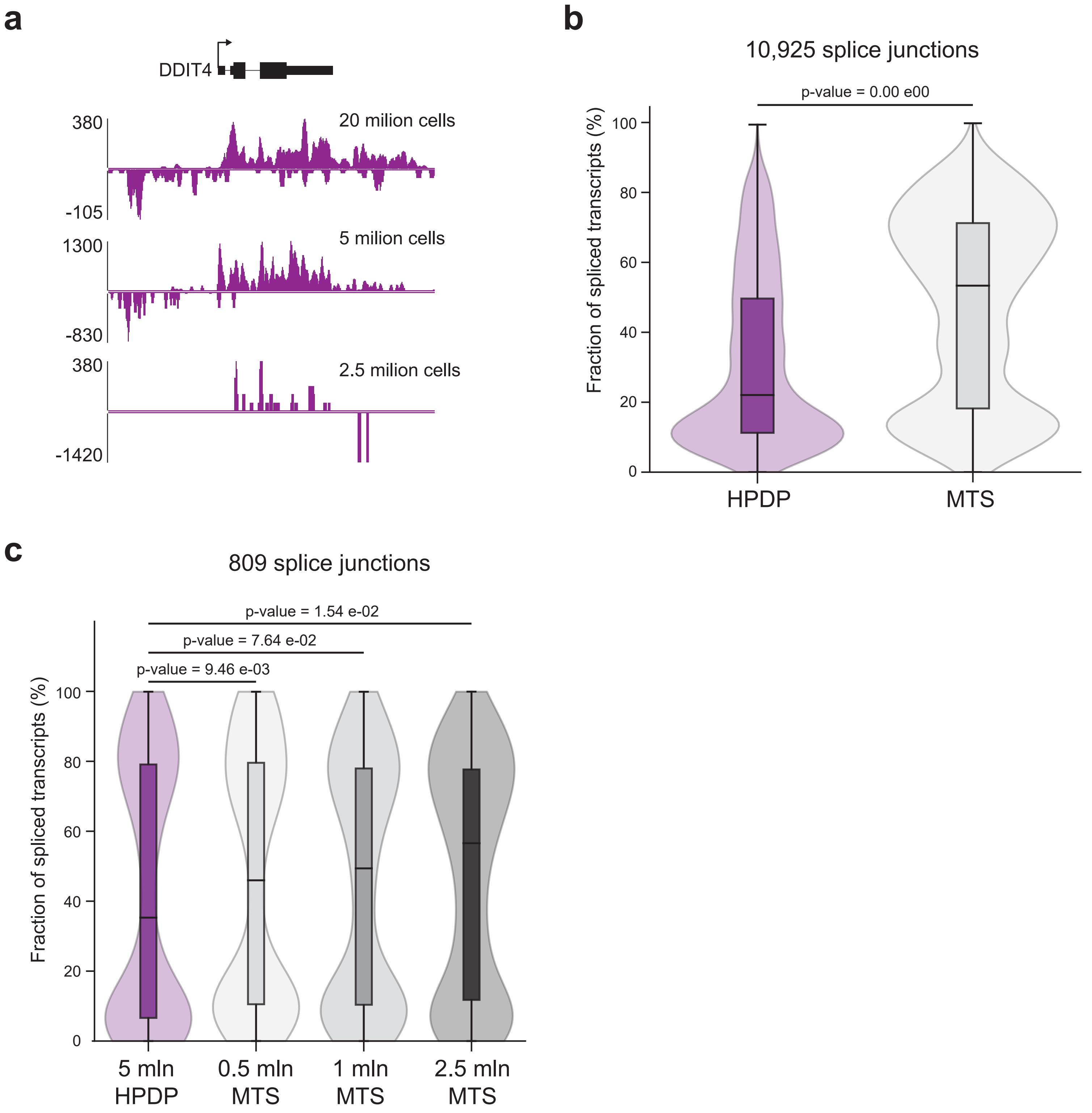
