## Supplementary Methods for "Rapid and scalable profiling of nascent RNA with fastGRO"

**fastGRO protocol** (Barbieri, Hill, Quesnel-Vallieres, Barash and Gardini, 2020)


### DAY ONE: Extraction of nuclei

Notes:

- cool centrifuge to 4* and add fresh Superase-In to buffers
- All stages done on ice or 4*.
- Collect 1 or 2 big plates of cells (15-30 mln total)
- Spike-in: isolate nuclei from Drosophila cells and add 5% of drosophila vs mammalian cells before NRO.

Buffers (add 2U/mL Superase-In before use):

- Swelling Buffer (500mL):
  - 500mL Ultrapure Water
  - 5mL 1M Tris-HCL pH 7.5 (final 10mM)
  - 1mL 1M MgCl2 (final 2mM)
  - 1.5ml 1M CaCl2 (final 3mM)
- Swelling Buffer + 10% Glycerol (200mL)
  - 180mL swelling buffer
  - 20mL glycerol
- Lysis Buffer (100mL)
  - 100mL Swelling buffer + 10% glycerol
  - 1mL Igepal (NP-40)
- Freezing Buffer (50mL)
  - 27.5mL Ultrapure water
  - 20mL of glycerol (final 40%)
  - 250ul of 1M MgCl2
  - 10ul of 0.5M EDTA (final 0.1mM)
  - 2.5mL Tris-HCL (use half pH 7.5, half pH8 so that at 4* the pH will be about 8.3) (final 50mM)

1. Wash cells with PBS.
2. Trypsinize as quick as possible.

3. Collect with cold medium in 50 ml falcon tube. Keep on ICE!
4. Wash twice with cold PBS.
5. Add 10mL of ice-cold swelling buffer to cells. Incubate for 5 minutes. Spin 400g for 10 minutes.
6. Remove supernatant and resuspend in 10mL Swelling Buffer with glycerol (should be at least 5x cell pellet)
7. Vortex lightly while adding equal volume of lysis buffer.
8. Let sit on ice for 5 minutes. Add 25mL lysis buffer and centrifuge 600g for 5 minutes.
9. Wash 1x: flick pellet to loosen and resuspend in 25mL lysis buffer. Centrifuge 600g for 5 minutes.
10. Remove supernatant and resuspend in 10mL freezing buffer. Take 10 µl for cell count.
11. Centrifuge 900g for 6 minutes and resuspend in freezing buffer to a concentration of 2x10^7 nuclei per 100 µL of freezing buffer (cut end of pipette tip to help resuspend viscous pellet).
12. Per library, use 1.5-2 x10^7 nuclei
13. Can store in -80 °C until ready to proceed.

**NRO reaction**

Buffers:

- 2x Nuclear run-on buffer (NRO). Will need 100ul/sample
    - 10 mM Tris-HCl pH8
    - 5 mM $MgCl_2$
    - 300 mM KCl
    - 1 mM DTT
    - 500 µM ATP
    - 500 µM GTP
    - 500 µM 4-thio-UTP
    - 2 µM CTP
    - 200 µ/ml Superase-in
    - 1% Sarkosyl (N-Laurylsarcosine sodium salt solution)
    - $H_2O$ to 100 µl/sample

1. Warm the NRO buffer at 30 °C.
2. Mix 100 µl of thawed nuclei solution (thawed on ice) with an equivalent volume of 2xNRO buffer. Pipette up and down 15 times using end-cut pipette tip.

3. Incubate 7 minutes at 30 °C.
4. Add 600 µl Trizol. Vortex. Incubate 5 minutes at RT.
    a. STOP POINT: Freeze with liquid nitrogen, and store at -80 °C.

**Total RNA precipitation**

1. Add 160 µl of chloroform, shake vigorously for 15s.
2. Incubate 2 minutes at RT.
3. Centrifuge at 4 °C and 12,000 rpm for 15 min.
4. Transfer upper, aqueous phase into new 1.5 ml centrifuge tube.
5. Add 400 µL of isopropanol to precipitate RNA.
6. Incubate at RT for 10 minutes.
7. Centrifuged at 4 °C and 12,000 rpm for 10 min.
8. Wash RNA pellet using 1 ml of cold 75 % ethanol and centrifuge at 4°C and 12,000 g for 10 min.
9. Completely remove ethanol and air-dry pellet.
10. Dissolve pellet in 100 µL of $H_2O$
11. Determine concentration by NanoDrop (or Qubit) spectrophotometer.

**RNA fragmentation via Bioruptor Plus**

1. Transfer 150-300 µg of RNA to a new 1.5 mL tube and add water up to 500 ul
    a. Save 5 µl as unfragmented RNA
    b. Add 5-10% of 4sU-labelled drosophila RNA if drosophila nuclei have not been used
    c. Settings: 1 cycle: 30 sec / 30 sec ON / OFF at high settings.
2. Transfer fragmented RNA to a 2 ml tube.
3. Analyze fragmentation efficiency running 1 µl of unfragmented and 1 µl of fragmented RNA on Agilent 2200 TapeStation or Bioanalyzer.
    a. STOP POINT: fragmented RNA can be frozen in liquid nitrogen and stored at -80 °C.

**DAY TWO: EZ-link HPDP-Biotinylation**

- Prepare 1 mg/ml EZ-link HPDP Biotin: resuspend 1 mg in 1 ml of DMF, vortex and incubate at 36 °C for 30 min. Store at -20 °C for long storage.

- 10x Biotinylation Buffer (BB) – prepare in advance and store at 4 °C:
    - 100 mM Tris pH 7.5
    - 10 mM EDTA pH 8.0

1. Heat samples at 65 °C for 10 minutes, incubate on ice for 5 minutes.
2. Add 100 µl of 10x BB, 200 µl of DMF and 200 µl of EZ-link HPDP Biotin in DMF.
3. Incubate in the dark at 24 °C and 800 rpm for 2 hours.

**Precipitation of biotinylated RNA**

1. Add 800 µl of chloroform to the RNA-biotin and mix.
2. Centrifuge at 15,000 rpm for 5 minutes.
3. Transfer upper phase (1 ml) into new tube.
4. Add 100 µl volume of 5 M NaCl and mix.
5. Add 1 ml of isopropanol and mix for 15 sec manually.
6. Centrifuge at 4 °C and 15,000 rpm for 30 min.
7. Remove supernatant.
8. Wash pellet with 1 mL of ice-cold 75 % ethanol.
9. Centrifuge at 4 °C and 15,000 rpm for 5 min.
10. Remove supernatant.
11. Quick spin at 4 °C and 15,000 rpm and remove remaining supernatant (<u>biotinylated RNA should not dry</u>).
12. Resuspend RNA in 100 µl $H_2O$
    a. STOP POINT: Store at -80 °C.

**Labeled RNA separation (Invitrogen Streptavidin Beads)**

- Wash Buffer (prepare freshly)
    - 100 mM Tris pH 7.5
    - 10 mM EDTA pH 8.0
    - 1M NaCl
    - 0.1% (v/v) Tween-20

1. Heat part of the WB at 65 °C (65°C-WB).
2. Prepare the beads:
    a. Take 100 µl of beads/sample.
    b. Wash the beads twice with 200 µl of wash buffer (2 Volumes)

    c. Resuspend in 1 Volume (100 µl/sample) of wash buffer

3. Add 100 µl of prepared Invitrogen streptavidin beads to 200 µl RNA-biotin.
4. Incubate at 4 °C in rotation for 15 min.
5. Transfer tubes containing RNA-bead-mix to a magnetic rack.
6. Wash 3-times with 900 µl of 65 °C-WB.
7. Wash 3-times with 900 µl of RT-WB.
8. Resuspend beads in in 100 µl of 100 mM DTT and incubate 5 minutes
9. Transfer tubes to the magnetic rack and collect the labeled RNA in a new tube.
10. Repeat the elution step and collect the eluted RNA in the same tube (200 µl final volume).

**RNA purification using RNA Clean and Purification kit-5 (Zymo Research) with DNAse step**

1. Use buffers provided with the Kit. Add ethanol to wash and pre-wash buffers and resuspend DNAse in water.
2. Add 2 volumes RNA Binding Buffer to each sample and mix (400 µl).
3. Add an equal volume of ethanol (95-100%) and mix (600 µl).
4. Transfer the sample to the Zymo-Spin IC Column in a Collection Tube and centrifuge for 30 seconds. Discard the flow-through.
5. Add 400 ul of RNA Wash Buffer to the column and centrifuge for 30 seconds at 16,000 g. Discard the flow-through.
6. For each sample to be treated, prepare DNase I reaction mix in an RNase-free tube. Mix well by gentle inversion:

   | | |
   |---|---|
   | DNase I | 5 µl |
   | DNA Digestion Buffer | 35 µl |

7. Add 40 µl reaction mix directly to the column matrix. Incubate at room temperature (20-30ºC) for 15 minutes.
8. Add 400 µl RNA Prep Buffer to the column and centrifuge for 30 seconds. Discard the flow-through.
9. Add 700µl RNA Wash Buffer to the column and centrifuge for 30 seconds. Discard the flow-through.
10. Add 400 µl RNA Wash Buffer to the column and centrifuge for 2 minutes. Discard the flow-through.

11. Centrifuge for 1 minute at full speed to ensure complete removal of the wash buffer. Transfer the column carefully into an RNase-free tube.
12. Add 6 µl DNase/RNase-Free Water directly to the column matrix, incubate for 1 minute and centrifuge for 30 seconds.
13. Measure eluted labelled RNA concentration by Qubit fluorometer.
14. Proceed to NEBNext Ultra II Directional RNA Library Prep Kit (NEB).
